# Unravelling the Molecular Footprints of Diabetic Foot Ulcers: In Silico Discovery of Key Protein and MicroRNA Signatures

**DOI:** 10.1101/2024.11.12.623130

**Authors:** Jay Mukesh Chudasama, Ghanshyam Parmar

**Affiliations:** Department of Pharmacy, Sumandeep Vidyapeeth Deemed to be University, Piparia, Waghodia, Vadodara 391760, Gujarat, India

**Keywords:** Diabetic Foot Ulcers (DFUs), Gene Ontology (GO) Analysis, MicroRNAs, hsa-miR-34a-5p, hsa-miR-155-5p, TP53, GAPDH, AKT1, MYC, TNF, EGFR, STAT3, FN1, VEGFA, JUN, Molecular Functions, Cellular Components, Pathways, Proteoglycans in Cancer, Human Cytomegalovirus Infection, Wound Healing, Inflammation

## Abstract

Diabetic foot ulcers (DFUs) present a significant clinical challenge, characterized by chronic inflammation and impaired wound healing. This study employs Gene Ontology (GO) analysis to identify critical biological processes, molecular functions, cellular components, and pathways associated with DFUs, aiming to uncover novel therapeutic targets. The analysis reveals significant enrichment in biological processes such as Positive Regulation of miRNA Transcription and Regulation of miRNA Transcription, highlighting the crucial role of microRNAs, including hsa-miR-34a-5p, hsa-miR-155-5p, hsa-miR-17-5p, hsa-miR-29b-3p, hsa-miR-7-5p, hsa-miR-1-3p, and hsa-miR-23b-3p, in regulating wound healing and inflammation. Enriched molecular functions, such as DNA-binding Transcription Activator Activity and Protein Phosphatase Binding, suggest that targeting genes like TP53, GAPDH, AKT1, MYC, TNF, EGFR, STAT3, FN1, VEGFA, and JUN could modulate critical cellular processes and improve DFU management. The analysis also identifies key cellular components, including Vesicle and Platelet Alpha Granule Lumen, as crucial for cellular transport and signaling, suggesting that interventions targeting these components could enhance wound repair. Furthermore, enriched pathways such as Proteoglycans in Cancer and Human Cytomegalovirus Infection indicate potential mechanisms and viral influences relevant to DFUs. These findings provide a comprehensive framework for developing targeted therapies that address the multifaceted pathology of DFUs, offering promising avenues for improving patient outcomes and advancing wound healing strategies.

## 1. Introduction

In recent years, diabetes mellitus has arisen as a major health issue on a global scale. This is because it is a chronic metabolic illness characterised by persistently high blood glucose levels. Diabetic foot ulcers (DFUs) are a particularly disabling complication of diabetes (1, 2). Chronic diabetic foot ulcers (DFUs) are most common in persons with diabetes and tend to form on the lower extremities. They are linked to a wide variety of risk factors, including neuropathy and poor circulation (3). Significant morbidity can ensue from these ulcers, diminishing patients’ quality of life while also raising healthcare expenses and, in extreme cases, necessitating amputation (4). Therefore, it is crucial for physicians and researchers to learn about the pathophysiology of DFUs, investigate the difficulties associated with treating them, and create new therapeutic approaches (5).

### Elaborating the Disease

Diabetes foot ulcers are a result of the complex interaction of metabolic abnormalities, neuropathy, and vascular alterations that frequently accompany diabetes mellitus. Diabetes neuropathy, a common consequence, affects peripheral nerves, resulting in sensory loss and motor impairment. Patients who have lost their protecting sense are more vulnerable to minor injuries, which can go unreported and escalate to ulcers over time. Concurrently, diabetes-related vascular alterations reduce blood flow to the extremities, decreasing the delivery of oxygen, nutrients, and immune cells required for wound healing. Furthermore, persistent hyperglycaemia promotes the accumulation of advanced glycation end products, which impair the normal function of wound-healing cells (6). As a result, diabetes patients reduced wound healing capacity allows for the formation of chronic, non-healing ulcers. DFUs manifest clinically in a variety of ways, ranging from superficial wounds to severe, infected lesions with underlying bone involvement (7). These ulcers are frequently classified based on their depth, size, and presence of infection, with such classifications assisting in the direction of treatment decisions. Despite advances in wound care and medical technology, treating DFUs remains difficult due to the complex underlying pathophysiology and the presence of several comorbidities that are common in diabetic patients (8).

### Common Deficits in the Study of Diabetic Foot Ulcers

Despite significant advances in our understanding of diabetic foot ulcers, there are still important gaps that prevent us from having a complete picture of this condition. The lack of universally accepted diagnostic criteria and evaluation methods for DFUs is a major shortcoming. Inconsistencies in patient care may result from differences in assessment methodologies, preventing reliable cross-study comparisons. For uniformity in research findings and to direct clinical decisions, it is critical to develop a standardised method of categorising DFUs (9, 10). Patient recruitment and retention are common problems in clinical trials investigating DFU management. Many diabetes patients have other comorbidities and adhere to complex care regimens, which might make participation in long-term trials difficult. This complicates efforts to collect reliable, long-term data that adequately represents the wide range of people living with diabetes (11). Furthermore, due to the possibility for adverse effects, performing placebo-controlled trials in this patient group requires novel trial designs that strike a balance between scientific rigour and patient safety. We also know very little about the molecular pathways that cause the delayed wound healing seen in DFUs, which is a significant gap. The particular molecular processes implicated in neuropathy and vascular insufficiency are still unknown. By better understanding these systems, we may be able to develop more effective therapeutic strategies for slow wound healing (12, 13).

### Novel work done by you

The prefund mode of action is explained in detail, making diabetic foot ulcer disease the primary focus of this study. With the help of this bioinformatics in silico study, researchers will have a solid foundation on which to build their hypotheses about the genes (proteins) and miRNAs that are linked to the complex diabetic foot ulcer disease and could be used as potential targets for treating it. All previously conducted and current research on diabetic foot ulcers is included in this study.

### In conclusion

Diabetic foot ulcers are a major obstacle in the treatment of diabetes and other wounds. Chronic, non-healing ulcers are a result of a complex interplay between neuropathy, vascular dysfunction, and metabolic abnormalities that can have devastating consequences. The lack of standardised evaluation methods, difficulties in clinical trial design, and an incomplete grasp of the underlying biological pathways all persist despite progress in research. Resolving these gaps is crucial for advancing our understanding of DFUs, boosting patient outcomes, and decreasing the prevalence of this disabling diabetic complication. Clinical practise and patient well-being stand to benefit greatly from the insights obtained by researchers doing this in silico investigation of diabetic foot ulcers (14).

## 2. Materials and Methods

### 2.1.Dataset Selection

Datasets related to Diabetic foot ulcer (DFU) were obtained from the Gene Expression Omnibus (GEO). The details about the selected diabetic foot ulcer related GSE datasets were elaborated below; **GSE134431** titled as Deregulated immune signature orchestrated by FOXM1 impairs human diabetic wound healing the experiment was performed through Expression profiling by high throughput sequencing; the comparison was done between the 8 diabetic foot skin and 13 diabetic foot ulcer (15). **GSE68183** titled with Comparative genomic, microRNA, and tissue analyses reveal subtle differences between non-diabetic and diabetic foot skin (gene expression); the study was carried out between 3 non-ulcerated non-neuropathic diabetic foot skin (DFS) and 3 healthy non-diabetic foot skin (NFS) (16). **GSE37265** titled with Transcriptome analysis of patients with recurrent aphthous stomatitis suggests novel therapeutic targets; this study was performed between 5 control and 14 ulcer patients. **GSE80178** titled with Genomic Profiling of Diabetic Foot Ulcers Identifies miR-15b-5p as a Major Regulator that Leads to Suboptimal Inflammatory Response and Diminished DNA Repair Mechanisms; and the study was performed between 3 non-diabetic foot skin and 9 diabetic foot ulcers.

### 2.2. Processing of Data and selection of Differentially Expressed Genes

The four datasets, GSE134431, GSE68183, GSE37265, and GSE80178, related to MS, were analyzed using the GEO2R tool (available at http://www.ncbi.nlm.nih.gov/geo/geo2r/) provided by GEO (https://www.ncbi.nlm.nih.gov/geo/). GEO2R generates a text file listing differentially expressed genes (DEGs) in disease versus control format, utilizing Bioconductor packages such as GEO query and limma R. This file includes LogFC values, p-values, adjusted p-values, gene symbols, gene IDs, and gene titles (17). DEGs with a p-value greater than 0.05 were excluded from the study, and genes with a fold change value (logFC) of at least 0.5 for both HD and MS were considered. After collecting data from the mentioned GSE datasets, we identified a large number of common or differentially expressed genes (DEGs). These genes were then filtered against Diabetic foot ulcer genes from GeneCard (https://www.genecards.org/Search/Keyword?queryString=diabetic%20foot%20ulcer), and the common genes were considered as differentially expressed genes (DGEs) (18).

### 2.3. Common DEG Identification and Network analysis

FunRich, a desktop tool, was used to identify the common DEGs in these two disorders. For gene network development, the online software GeneMANIA was employed. The common DEGs were imported into STRING (Search Tool for the Retrieval of Interacting Genes) (https://string-db.org/), and a protein-protein interaction (PPI) network was created using STRING’s database (https://string-db.org/cgi/input?sessionId=bU53xJ1FawGP&input_page_active_form=multiple_identifiers). This PPI network was then directly imported into Cytoscape from STRING. Using the Cytoscape plugin CytoHubba, the gene in the DEGs list with the most connections to other genes (hub genes) was identified. Enrichment analysis was performed using the online software Enrichr. The DEGs with the highest connectivity in the PPI were further analyzed for microRNA interactions using the online software miRNet (19–21).

## 3. Results

In this present study, we used the microarray data available from Gene Expression Omnibus for Diabetic foot ulcer (DFU) disease. We studied the differentially expressed genes (DGEs), nature of protein-protein interactions, identified the hub genes, and also identified the mi RNAs associated with the top 10 hub genes (22). The data used for this study has been normalized (**Fig. 1**).

**Fig. 1:**
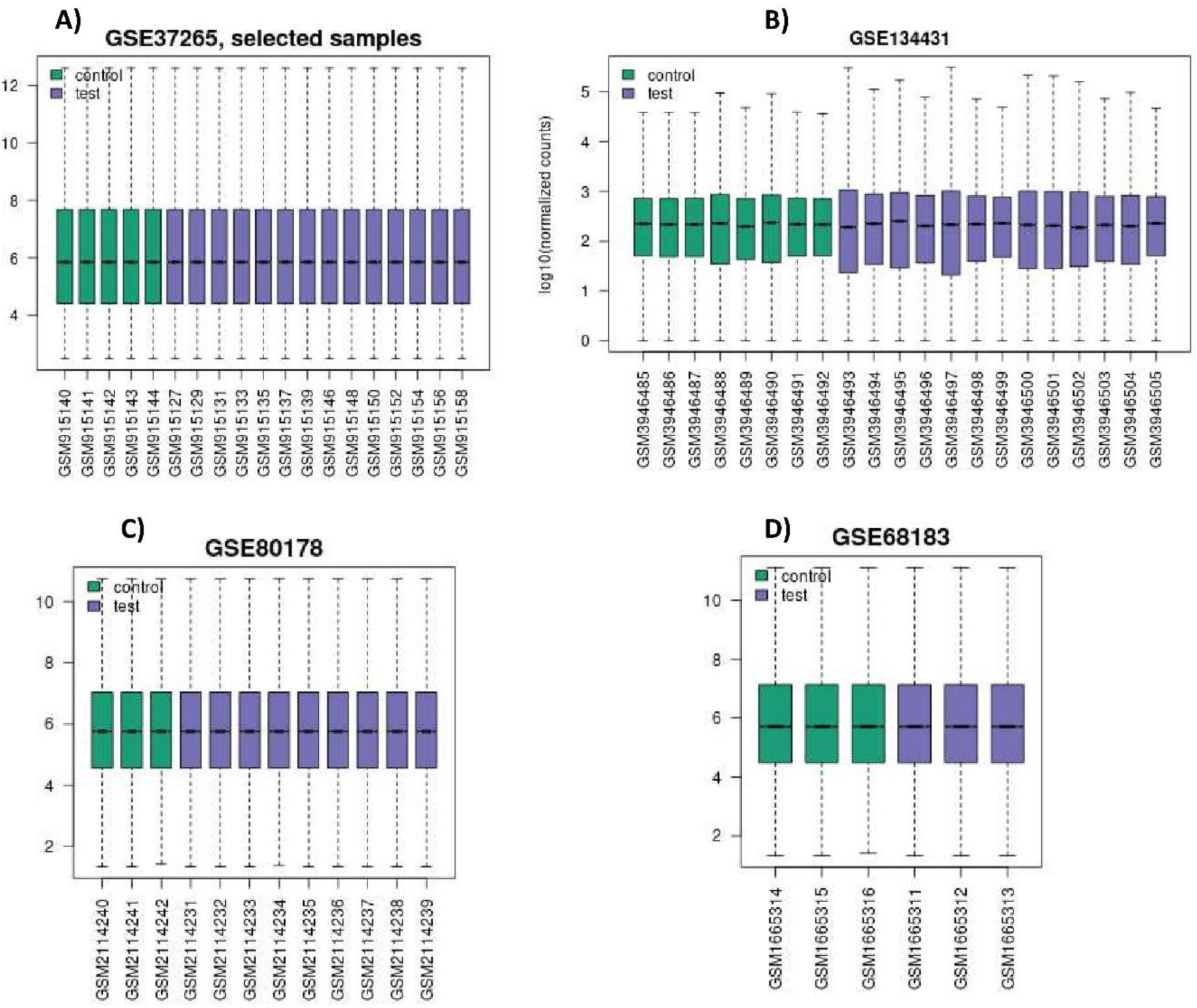
Box plots of DFU representing the normalization of data. The distribution of Control versus Diseased samples was depicted after data normalization with the GEO2R program. Where A) GSE37265, B) GSE134431, C) GSE80178, D) GSE68183. Each box plot represents the differential gene expression value for single patients.

### 3.1. Finding the common DEGs in these datasets

The DEGs shared among the GSE datasets were found to total 10,031, as shown in Figure 2. Due to the large number of genes and their complexity, these common genes from the GSE datasets were further filtered using the genes obtained from GeneCard. A Venn diagram was then created to compare the common genes from the GSE datasets with the GeneCard genes, revealing that 1,435 genes were shared between the two sets, as depicted in Figure 3. These 1,435 common genes were subsequently selected for further studies (23, 24).

**Fig. 2:**
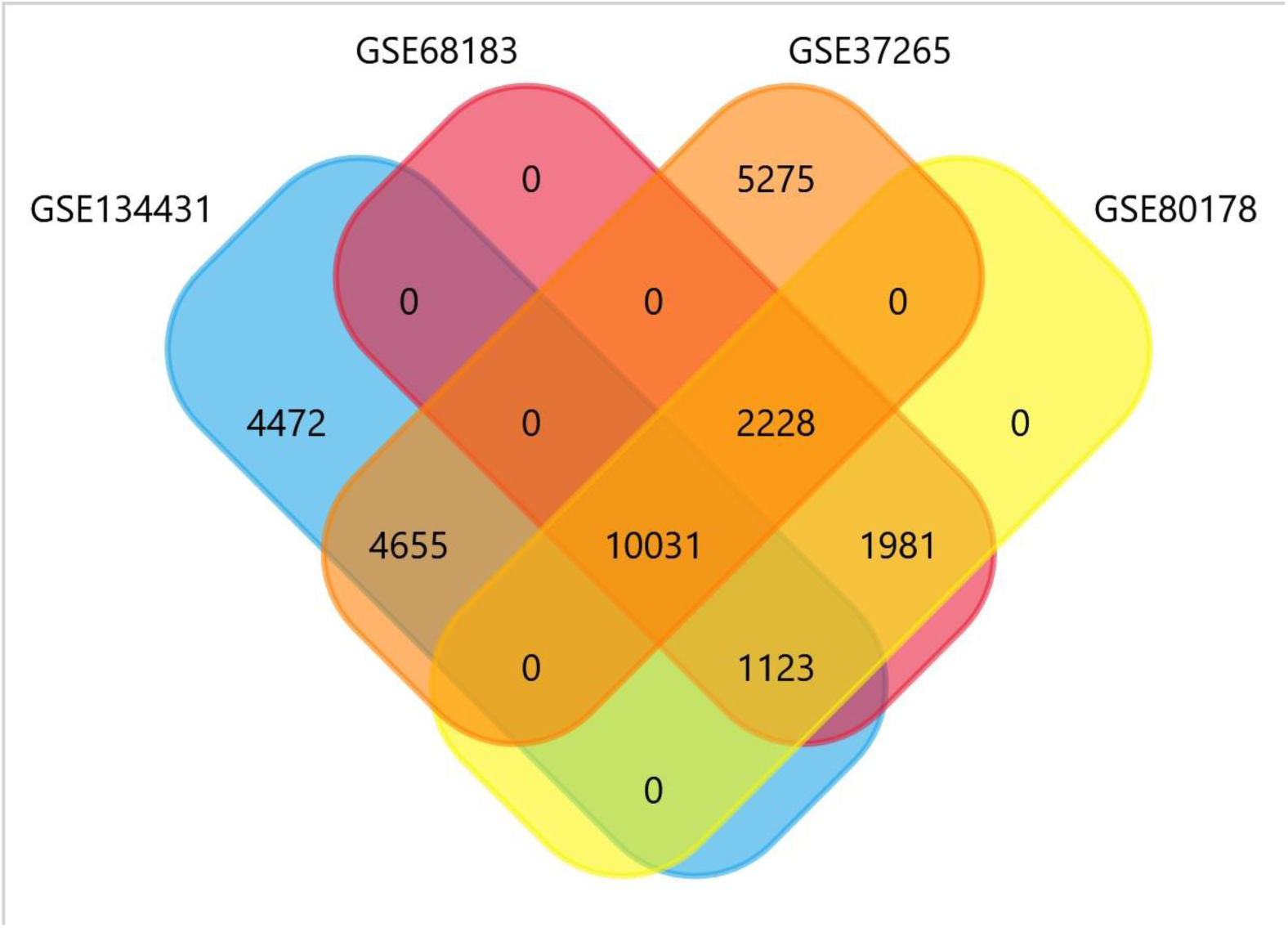
Venn diagram which represents the number of common differentially expressed genes between various GSE datasets (GSE37265, GSE68183, GSE80178, and GSE13443)

**Fig. 3:**
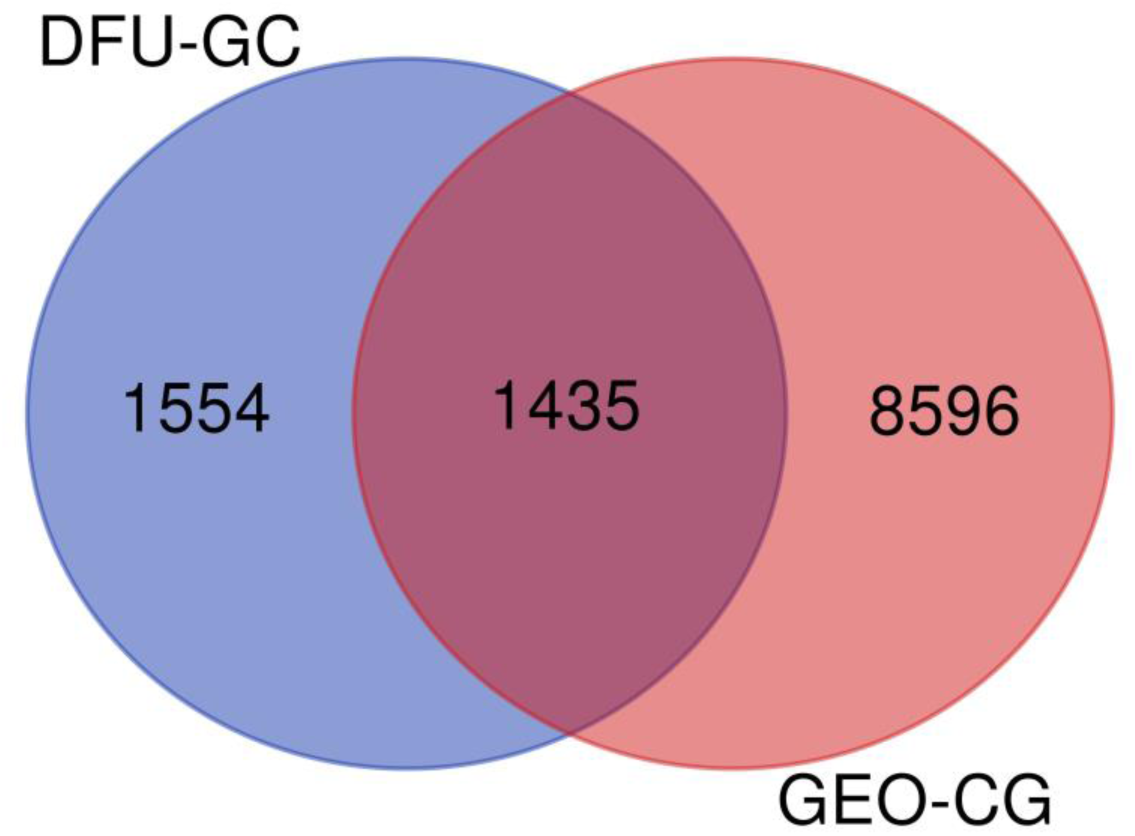
Venn diagram which represents the number of common differentially expressed genes between the common DEGs of GSE datasets (GEO-CG) and DFU GeneCard (DFU-GC)

### 3.2. Interaction between common DEGs

The interactions among the common DEGs were visualized using STRING. The list of common DEGs was input into the STRING database, which employs the Markov Clustering Algorithm to develop a protein-protein interaction (PPI) network (Figure 4). This resulted in a dense gene network for the DEGs.

**Fig. 4:**
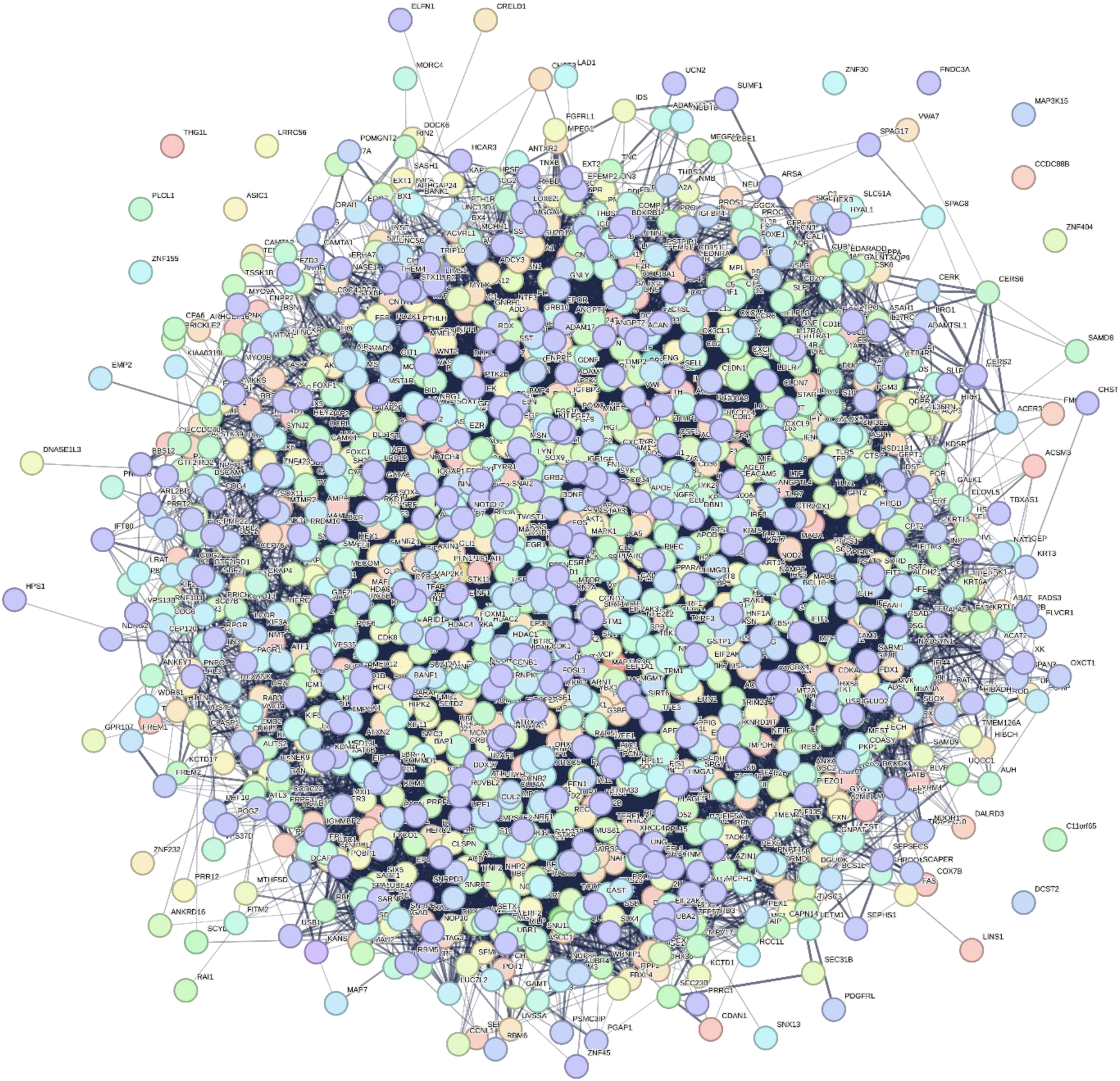
The protein-protein interaction network was developed by using the MCL clustering method. For this network, DFU GeneCard genes and GSE datasets DEGs were used. In this network, the interaction score was set at medium confidence (0.400).

Within the PPI, some proteins did not interact at all, while others had intriguing connections to multiple proteins in the network. To identify hub genes, the PPI was imported into Cytoscape software, and the CytoHubba plugin was used to visualize and calculate the degree of centrality for all the hub genes (Table 1) (25).

**Table 1:** Top ten Hub proteins in the PPI network obtained from the DEGs of the datasets for HD and MS with their degree of centrality.

| Rank | Name of the protein | Degree score |
| --- | --- | --- |
| 1 | TP53 | 483 |
| 2 | GAPDH | 472 |
| 3 | AKT1 | 440 |
| 4 | MYC | 398 |
| 5 | TNF | 383 |
| 6 | EGFR | 363 |
| 7 | STAT3 | 321 |
| 8 | FN1 | 302 |
| 9 | VEGFA | 301 |
| 10 | JUN | 297 |

The top ten hub genes were selected for constructing a PPI network (Figure 5). Cellular tumor antigen p53 (TP53) had the highest degree of centrality (degree 483), followed by Glyceraldehyde-3-phosphate dehydrogenase (GAPDH) with a degree of 472, and RAC-alpha serine/threonine-protein kinase (AKT1) with a degree of 440. Myc proto-oncogene protein (MYC) had a degree of 398, Tumor necrosis factor (TNF) had a degree of 383, Epidermal growth factor receptor (EGFR) had a degree of 363, Signal transducer and activator of transcription 3 (STAT3) had a degree of 321, Fibronectin 1 (FN1) had a degree of 302, Vascular endothelial growth factor A (VEGFA) had a degree of 301, and Transcription factor Jun (JUN) had the lowest degree (297). All these genes were visualized clearly when the interaction score in STRING was set to high confidence (0.700) (26).

**Fig. 5:**
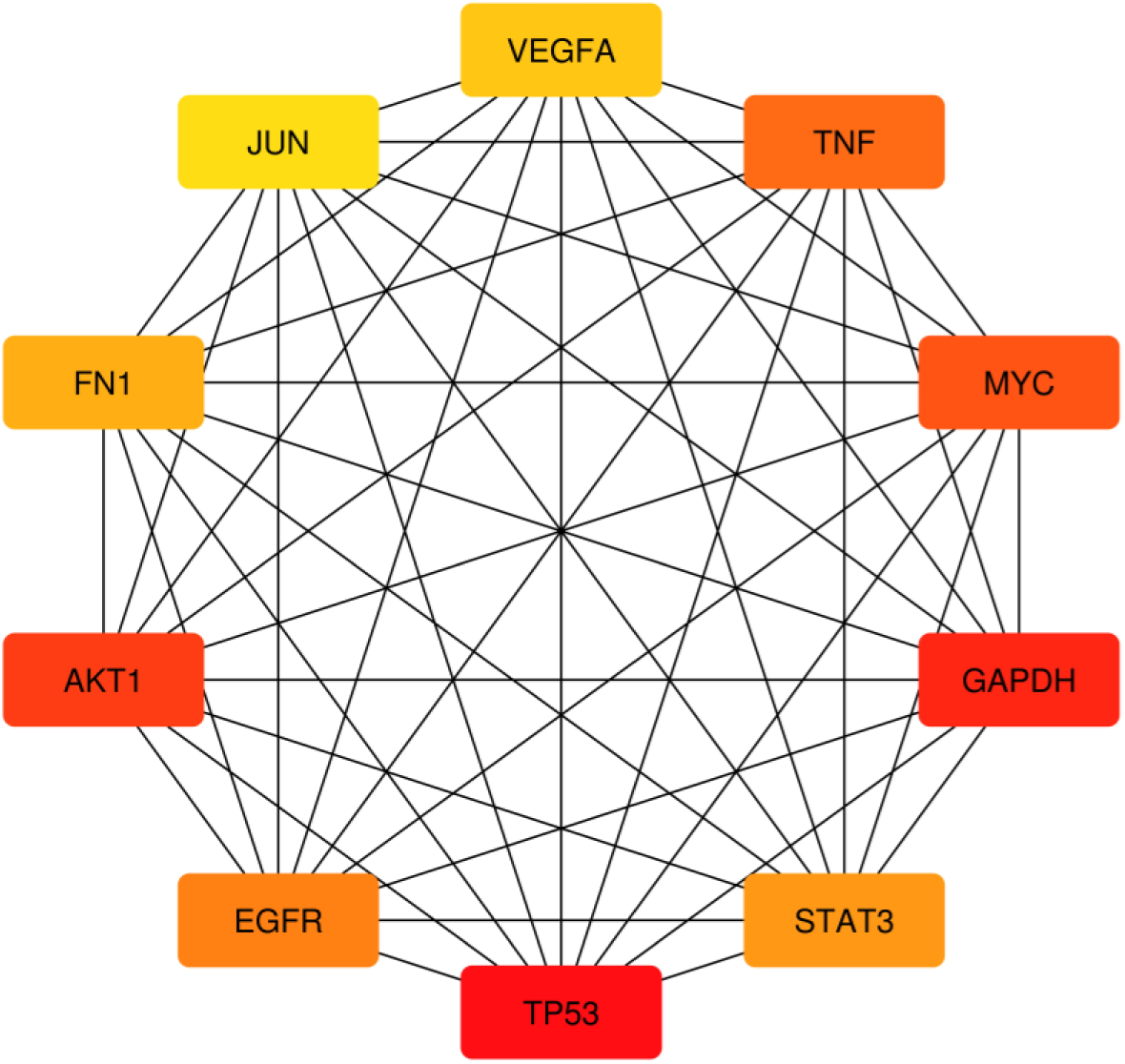
Network of top ten hub genes which was obtained using Cytoscape. The colour code represents the degree here, the red colour denotes the highest degree of centrality, orange denotes medium degree and yellow denotes the lowest degree of centrality.

PPI network showing the top ten hub genes was then obtained by setting the interaction score to high confidence (0.700) (**Fig. 6**).

**Fig. 6:**
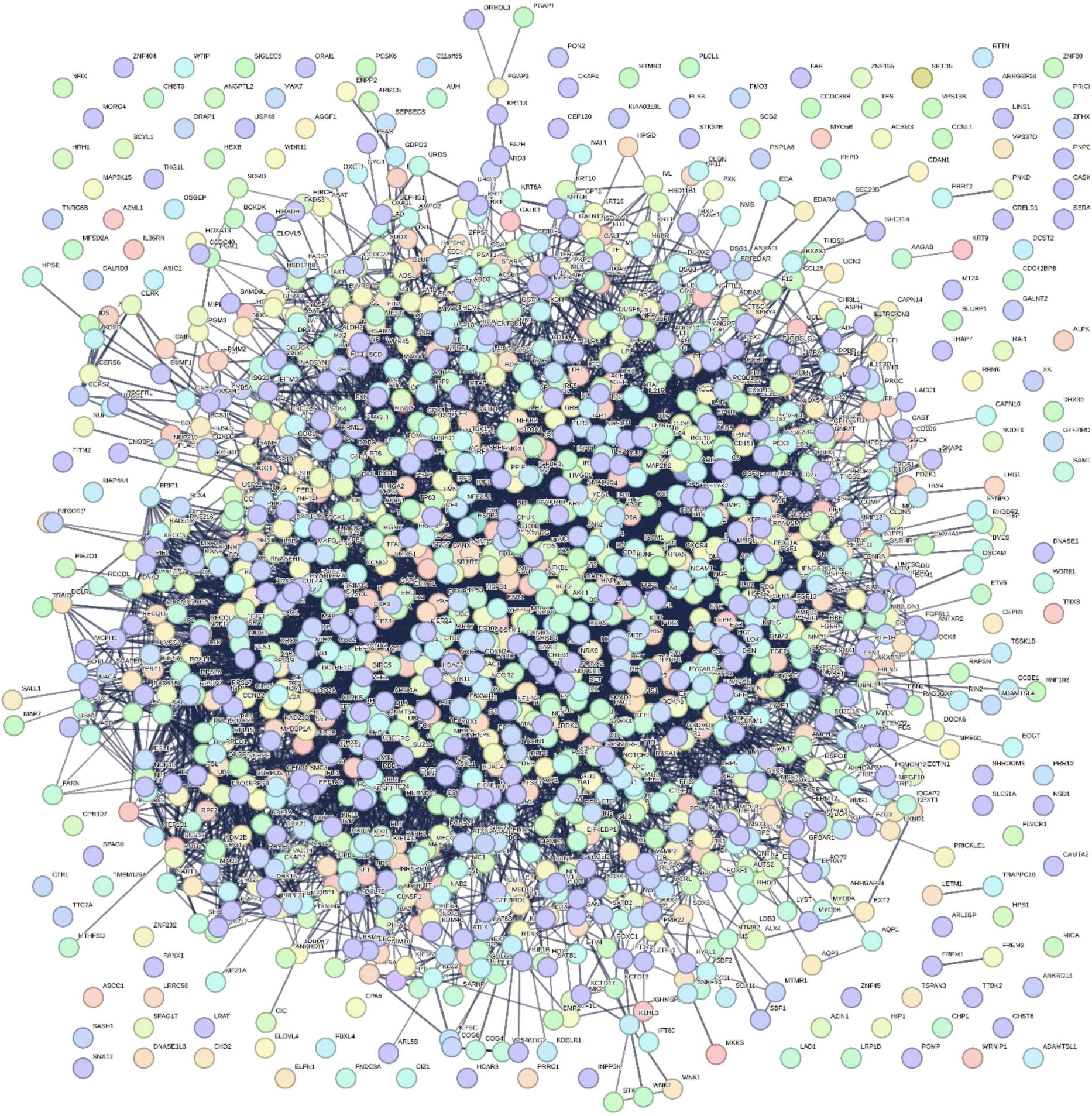
The PPI network showing the hub genes. This particular network was developed by setting the interaction score to high confidence (0.700), a high confidence score was used to reduce the interactions and to clearly state the Hub genes interacting with other genes.

### 3.3. Identification of miRNA and transcription factors associated with DEGs

The top ten hub genes were submitted to miRNet, an online platform to identify gene-miRNA interactions. This generated a network illustrating the interactions between DEGs, miRNAs, and transcription factors (Figure 7). The analysis revealed 251 miRNAs and fifty-eight transcription factors associated with the ten DEGs. In this network, MYC had the highest connectivity (degree 168), followed by VEGFA (degree 144). The miRNAs hsa-mir-34a-5p and hsa-mir-155-5p showed the highest connectivity (degree 10), followed by hsa-mir-17-5p, hsa-mir-29b-3p, hsa-mir-7-5p, hsa-mir-1-3p, and hsa-mir-23b-3p, each with a degree of 8. The miRNAs and transcription factors associated with these ten DEGs are listed in Table 2 (27, 28).

**Fig. 7:**
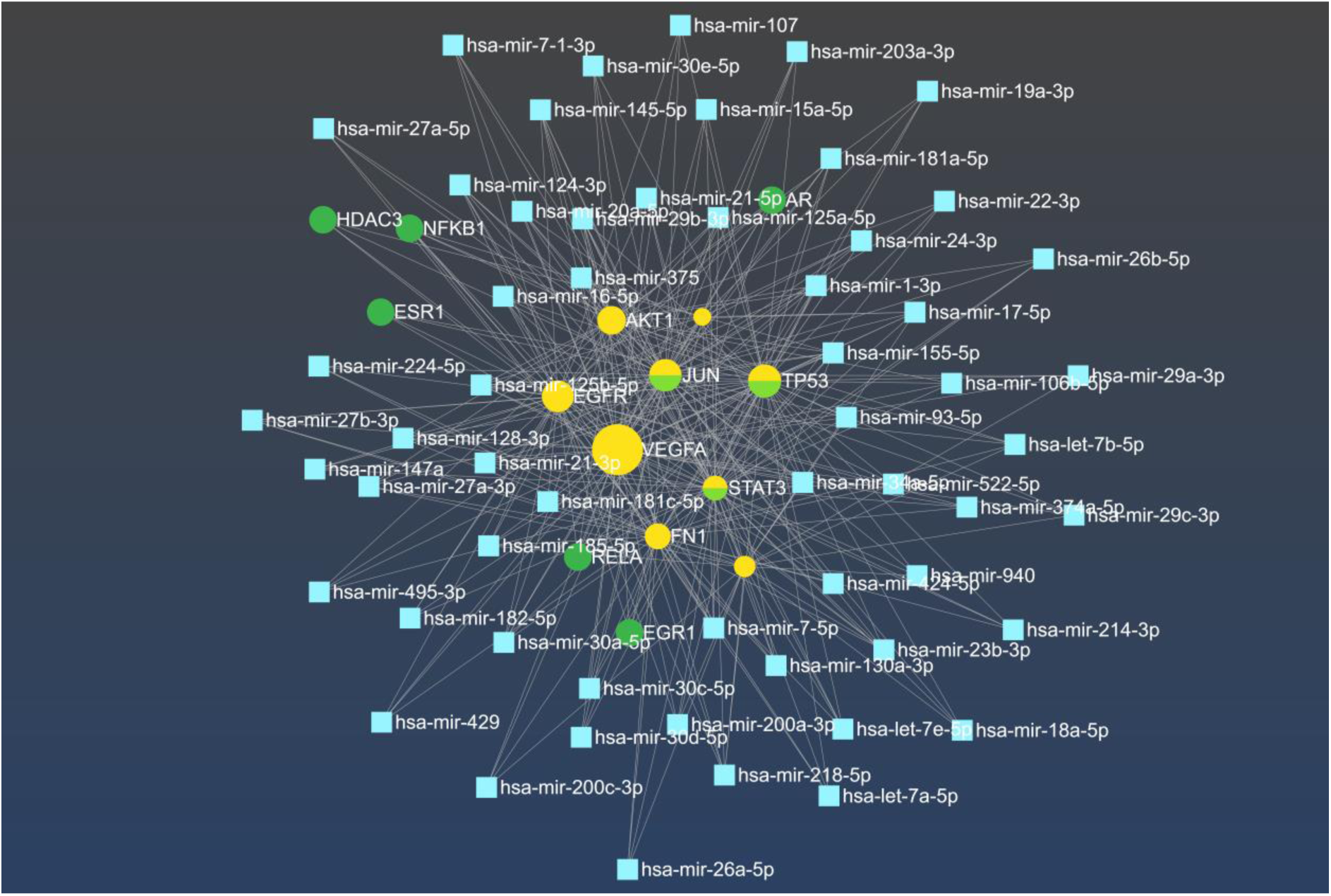
The network shows the interaction among DEGs, miRNAs, and transcription factors generated using miRNet. The genes are represented in yellow, the miRNA is represented in blue, transcription factors in green and transcription factors with identical gene and protein names are dual-colored (i.e. green and yellow-colored). The genes MYC and VEGFA have maximum connectivity (degree 168 and 144 respectively) and the miRNA hsa-mir-155-5p and hsa-mir-34a-5p are the ones with maximum connectivity (both have degree 10).

**Table 2:** List of miRNAs and transcription factors together with 10 DEGs with their degree of centrality common to Diabetic foot ulcer.

| Names of DEGs, miRNA, and transcription factors | Degree |
| --- | --- |
| MYC | 168 |
| VEGFA | 144 |
| JUN | 127 |
| TP53 | 127 |
| STAT3 | 102 |
| FN1 | 90 |
| EGFR | 89 |
| AKT1 | 80 |
| GAPDH | 60 |
| TNF | 25 |
| hsa-mir-34a-5p | 10 |
| hsa-mir-155-5p | 10 |

### 3.4. Common GO processes and KEGG pathways for the DEGs associated with HD and MS

For the enrichment analysis, the DEGs were submitted to the online software, Enrichr, and GO biological processes, GO molecular function, GO cellular component and KEGG pathways options were used for the enrichment analysis.

#### 3.4.1. GO biological processes enriched which are common

The biological process that is significantly most enriched is the Positive Regulation Of miRNA Transcription (GO:1902895) followed by the Positive Regulation Of miRNA Metabolic Process (GO:2000630) and Regulation Of miRNA Transcription (GO:1902893). The graphical representation of the GO biological processes common to the set of DEGs submitted has been represented graphically (**Fig 8a**) (29).

**Fig. 8:**
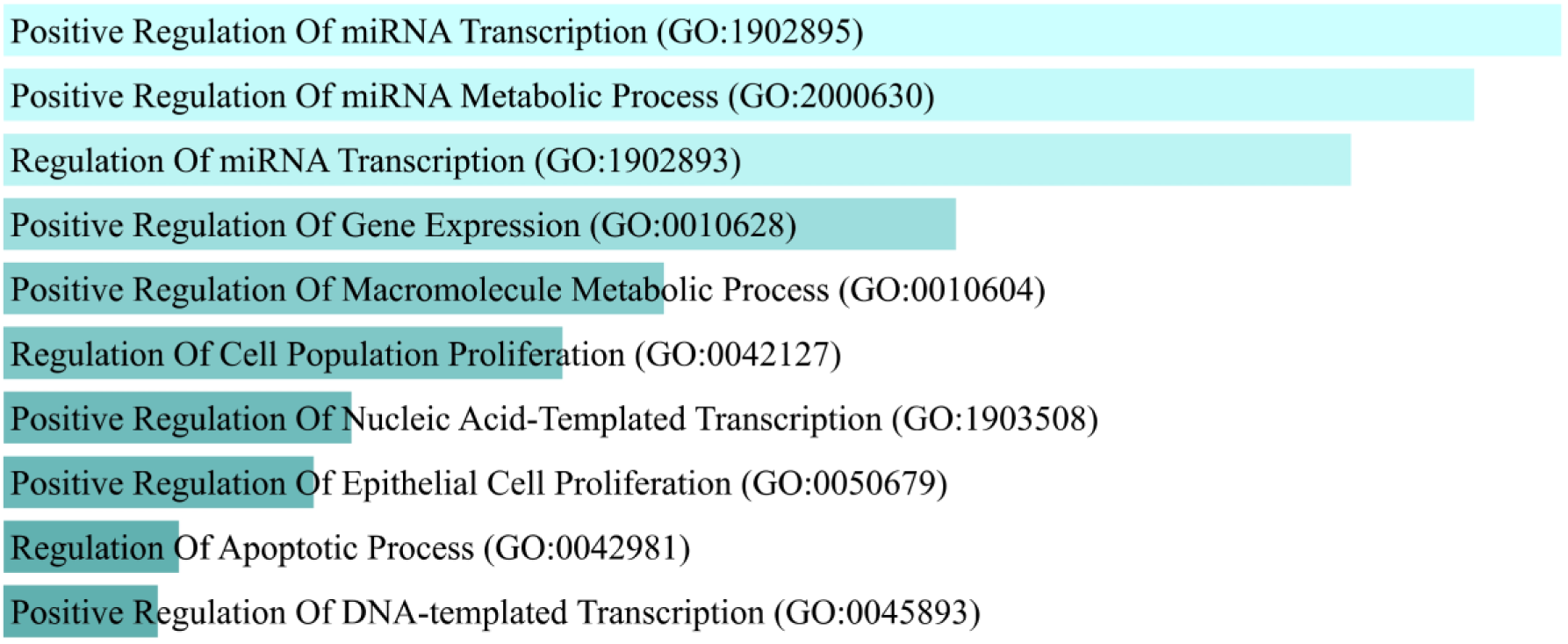
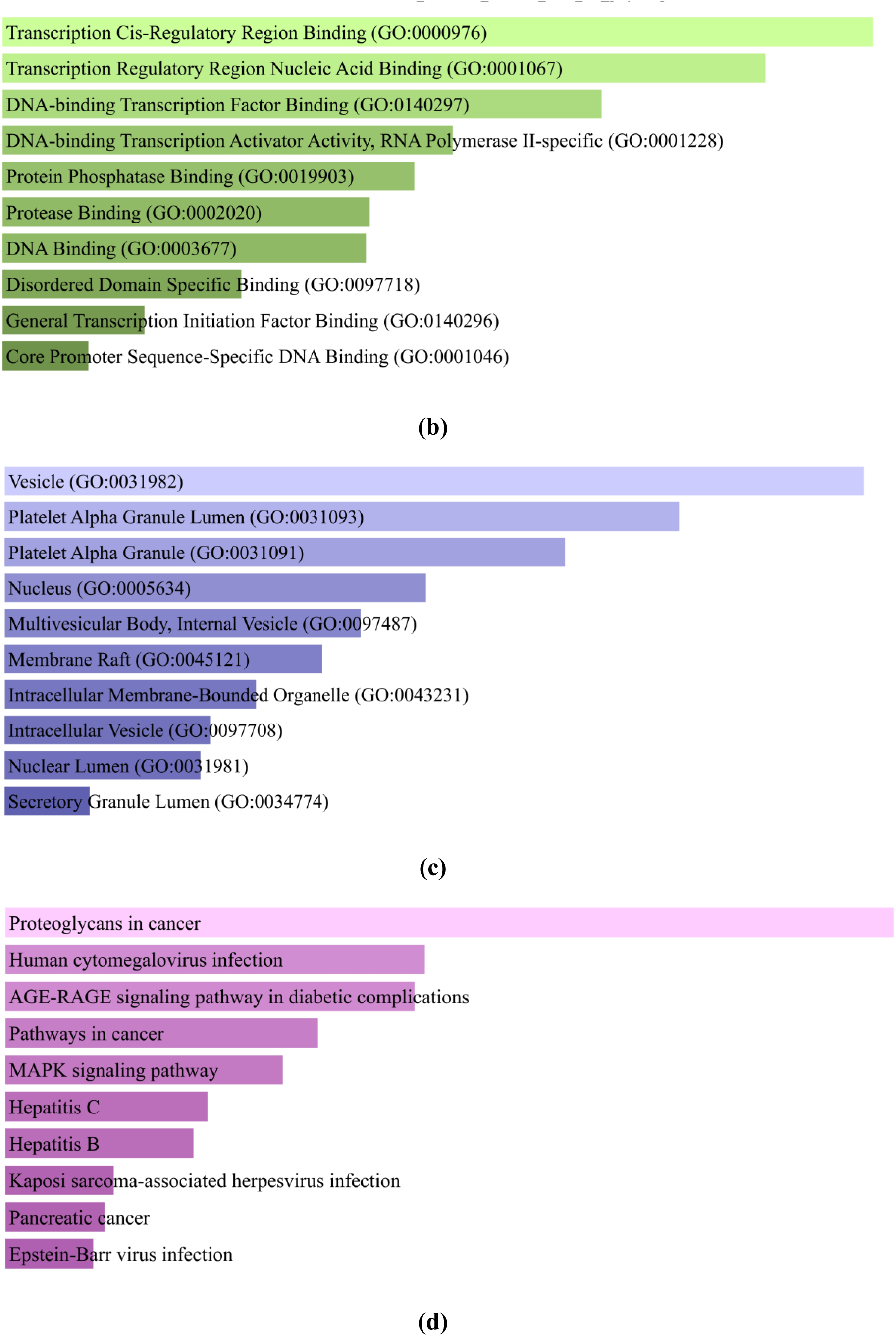
The enrichment analysis results in a graphical format. **(a)** The top ten biological processes enriched in the chosen set of DEGs. The x-axis represents the number of genes and the y-axis represents biological processes. **(b)** Top ten molecular functions that are enriched in these datasets. The x-axis represents the number of genes and the y-axis represents the molecular functions. **(c)** The top 10 cellular components are enriched in the DEGs of interest. The x-axis represents the number of genes and the y-axis represents the cellular components. **(d)** The DEGs of interest are enriched in the top 10 cellular components. The x-axis represents the number of genes and the y-axis represents the enriched KEGG pathways.

#### 3.4.2. Common GO molecular functions enriched

The Transcription Cis-Regulatory Region Binding (GO:0000976) with a p-value of 0.000001672 followed by Transcription Regulatory Region Nucleic Acid Binding (GO:0001067) with a p-value of 0.000003051 and DNA-binding Transcription Factor Binding (GO:0140297) with a p-value of 0.000007600. The molecular functions which are enriched with these are DNA-binding Transcription Activator Activity, RNA Polymerase II-specific, Protein Phosphatase Binding, Protease Binding, DNA Binding, Disordered Domain Specific Binding, General Transcription Initiation Factor Binding, General Transcription Initiation Factor Binding. The graphical representation of the enriched molecular functions has been represented in (**Fig 8b**) (30).

#### 3.4.3. Common GO cellular components enriched

The cellular components which are enriched are the Vesicle (GO:0031982) with a p-value of 0.0001808 followed by the Platelet Alpha Granule Lumen (GO:0031093) with a p-value of 0.0004745 and Platelet Alpha Granule (GO:0031091) with a p-value of 0.0008609. The other cellular components include Nucleus, Multivesicular Body-Internal Vesicle, Membrane Raft, Intracellular Membrane-Bounded Organelle, Intracellular Membrane-Bounded Organelle, Nuclear Lumen, Secretory Granule Lumen. (**Fig 8c**) illustrates the graphical display of enhanced cellular components (31).

#### 3.4.4. Common KEGG pathways that are enriched

The pathways named Proteoglycans in cancer with a p-value of 4.696e-15, followed by Human cytomegalovirus infection (p-value of 2.422e-12), Human cytomegalovirus infection (p-value of 2.772e-12), Pathways in cancer, MAPK signaling pathway, Hepatitis C, Hepatitis B, Kaposi sarcoma-associated herpesvirus infection, Pancreatic cancer, Epstein-Barr virus infection. (**Fig 8d**) shows a graphical representation of enriched cellular components.

## 4. Discussion

Diabetic foot ulcers (DFUs) are a serious complication of diabetes, characterized by chronic wounds on the feet that are difficult to heal due to a combination of factors such as neuropathy, peripheral arterial disease, and immune dysfunction, all of which are worsened by hyperglycemia. These ulcers can lead to severe outcomes, including infections, gangrene, and potential amputation if not managed properly. The pathogenesis of DFUs involves the dysregulation of key genes, including TP53, GAPDH, AKT1, MYC, TNF, EGFR, STAT3, FN1, VEGFA, and JUN, which are critical in processes such as inflammation, angiogenesis, and cellular repair. Targeting these genes offers the potential to correct the disrupted pathways, thereby enhancing wound healing and reducing inflammation. MicroRNAs (miRNAs), which are small non-coding RNAs that regulate gene expression by binding to the mRNA of target genes, have shown promise as therapeutic tools in this context. Specifically, hsa-miR-155-5p and hsa-miR-34a-5p have been identified as having significant connectivity with differentially expressed genes (DEGs) in DFUs. hsa-miR-155-5p is involved in regulating inflammatory responses and has been linked to the chronic inflammation seen in DFUs. By targeting genes such as TNF and STAT3, hsa-miR-155-5p can potentially reduce inflammation and promote a more balanced immune response, aiding in wound healing. Similarly, hsa-miR-34a-5p is associated with cell cycle regulation, apoptosis, and angiogenesis, targeting genes like TP53 and VEGFA. Through these interactions, hsa-miR-34a-5p can influence cell proliferation and new blood vessel formation, both of which are crucial for effective wound healing in DFUs. By modulating these miRNAs, there is potential to restore normal gene expression patterns, promote wound healing, and prevent the chronic progression of DFUs (32, 33).

The datasets related to diabetic foot ulcers (DFUs) were carefully gathered from the Gene Expression Omnibus (GEO), a comprehensive public repository that stores high-throughput gene expression data. These datasets, identified by their unique GEO Series (GSE) numbers, provide valuable insights into the molecular mechanisms underlying DFUs and form a crucial foundation for identifying novel therapeutic targets. Among these datasets, GSE134431, titled “Deregulated immune signature orchestrated by FOXM1 impairs human diabetic wound healing,” focuses on understanding immune dysregulation in diabetic wound healing. This study, conducted using expression profiling by high-throughput sequencing, compared gene expression profiles between 8 samples of diabetic foot skin and 13 samples of diabetic foot ulcers, revealing the critical role of the FOXM1 transcription factor in orchestrating an impaired immune response in diabetic wounds. This dataset is vital for exploring how immune system dysregulation contributes to poor healing outcomes seen in DFUs. Another significant dataset, GSE68183, titled “Comparative genomic, microRNA, and tissue analyses reveal subtle differences between non-diabetic and diabetic foot skin (gene expression),” investigates the genomic differences between diabetic and non-diabetic foot skin. The study analyzed 3 samples of non-ulcerated, non-neuropathic diabetic foot skin (DFS) and 3 samples of healthy non-diabetic foot skin (NFS), providing insights into early molecular changes in diabetic skin before ulcer formation. The findings from this dataset can aid in identifying biomarkers for early detection and prevention of DFUs (34). Additionally, GSE37265, titled “Transcriptome analysis of patients with recurrent aphthous stomatitis suggests novel therapeutic targets,” although primarily focusing on recurrent aphthous stomatitis, offers valuable insights into ulcerative conditions, including DFUs. This study, involving the transcriptome analysis of 5 control patients and 14 ulcer patients, provides a broad view of gene expression changes associated with ulcer formation and healing, which can be cross-referenced with DFU-related datasets to identify common pathways and potential therapeutic targets for ulcer management. The GSE80178 dataset, titled “Genomic profiling of diabetic foot ulcers identifies miR-15b-5p as a major regulator that leads to suboptimal inflammatory response and diminished DNA repair mechanisms,” highlights the role of microRNAs in DFU pathology. Conducted on 3 samples of non-diabetic foot skin and 9 samples of diabetic foot ulcers, this analysis identified miR-15b-5p as a key regulator of the inflammatory response and DNA repair mechanisms in DFUs, pointing to its potential as a therapeutic target. This dataset is particularly important for understanding the post-transcriptional regulation of genes involved in DFU pathogenesis. These GEO datasets are invaluable for researchers seeking to uncover the molecular underpinnings of DFUs (35). By providing access to raw gene expression data, they enable the identification of differentially expressed genes (DEGs) and pathways crucial to the development and progression of DFUs. Researchers can perform integrative analyses, such as gene set enrichment analysis (GSEA), to identify novel therapeutic targets and biomarkers. The availability of both mRNA and microRNA expression data also allows for the study of complex regulatory networks governing gene expression in DFUs. For example, datasets like GSE80178, which focus on microRNAs, offer insights into how these small non-coding RNAs modulate key genes involved in inflammation, angiogenesis, and wound healing, paving the way for new therapeutic strategies to improve healing outcomes in DFU patients. The genes identified in these datasets, including those involved in immune response, inflammation, angiogenesis, and DNA repair, represent potential targets for novel DFU therapies. By analyzing the expression patterns of these genes in diabetic and non-diabetic tissues, researchers can pinpoint key drivers of DFU pathology. For instance, identifying upregulated genes in ulcerated tissues compared to non-ulcerated tissues can highlight therapeutic targets aimed at reducing inflammation or enhancing tissue repair. Moreover, datasets like GSE134431 and GSE80178, which focus on immune dysregulation and microRNA regulation, provide a deeper understanding of how specific genes and pathways contribute to the chronic nature of DFUs. Targeting these pathways with gene therapy, small molecules, or biologics could offer new avenues for treatment, ultimately improving the quality of life for patients with diabetic foot ulcers (36).

By constructing a Venn diagram of the previously mentioned datasets, we identified 10,031 genes that were common across the selected GSE datasets related to diabetic foot ulcers (DFUs). To refine this extensive list, a further comparison was made using GeneCards, a comprehensive database of human genes, specifically focusing on genes associated with DFU. This process involved forming another Venn diagram to determine the overlap between the common genes from the GSE datasets and the genes listed in GeneCards. As a result, 1,435 genes were identified as the most effective differentially expressed genes (DEGs) that could be targeted for DFU treatment. These 1,435 DEGs represent the most promising candidates for further research and potential therapeutic intervention in DFUs (37, 38).

The identification of the top ten hub genes in the protein-protein interaction (PPI) network is a significant achievement in understanding the molecular mechanisms underlying diabetic foot ulcers (DFUs). These hub genes, characterized by their high degree of connectivity, represent critical nodes in the network that play essential roles in cellular processes and disease progression. The Cellular tumor antigen p53 (TP53), with the highest degree of 483, is a well-known tumor suppressor gene involved in regulating the cell cycle, apoptosis, and DNA repair, making it a key player in wound healing and tissue regeneration. Glyceraldehyde-3-phosphate dehydrogenase (GAPDH), with a degree of 472, is not only a central enzyme in glycolysis but also has roles in gene regulation and apoptosis, highlighting its importance in energy metabolism and cellular homeostasis in DFUs. RAC-alpha serine/threonine-protein kinase (AKT1), with a degree of 440, is involved in various cellular processes, including cell survival, proliferation, and metabolism, making it crucial for maintaining cellular integrity in the face of chronic wounds. Myc proto-oncogene protein (MYC), with a degree of 398, is a transcription factor that regulates the expression of numerous genes involved in cell growth and proliferation, underscoring its role in tissue repair and regeneration (39, 40). Tumor necrosis factor (TNF), with a degree of 383, is a key mediator of inflammation, which is a central feature of DFUs, and its regulation is crucial for controlling chronic inflammation in wounds. Epidermal growth factor receptor (EGFR), with a degree of 363, is involved in cell growth and differentiation and is critical for re-epithelialization during wound healing. Signal transducer and activator of transcription 3 (STAT3), with a degree of 321, is a transcription factor that mediates the expression of various genes involved in cell growth, survival, and immune response, making it a vital player in wound healing. Fibronectin 1 (FN1), with a degree of 302, is a glycoprotein that plays a crucial role in cell adhesion, migration, and tissue repair, highlighting its importance in the wound healing process. Vascular endothelial growth factor A (VEGFA), with a degree of 301, is a key regulator of angiogenesis, the formation of new blood vessels, which is essential for supplying nutrients and oxygen to healing tissues. Lastly, the transcription factor Jun (JUN), with a degree of 297, is involved in regulating gene expression in response to stress and is crucial for cellular response to injury (41). The analysis of microRNA (miRNA) interactions with these DEGs further emphasizes their regulatory significance. The submission of these top ten hub genes to miRNet, an online platform for gene-miRNA interaction analysis, revealed a complex network involving 251 miRNAs and fifty-eight transcription factors associated with the DEGs. In this network, the MYC gene emerged as the one with maximum connectivity (degree 168), followed by VEGFA (degree 144), indicating their central roles in the regulatory network of DFUs (42).

Among the miRNAs, hsa-miR-34a-5p and hsa-miR-155-5p both exhibited the highest connectivity in the network (degree 10), suggesting their significant roles in modulating the expression of these hub genes. These miRNAs are known to be involved in key processes such as apoptosis, inflammation, and cellular stress responses, making them critical regulators in the context of DFUs. Other miRNAs, including hsa-miR-17-5p, hsa-miR-29b-3p, hsa-miR-7-5p, hsa-miR-1-3p, and hsa-miR-23b-3p, also showed substantial connectivity (degree 8), further highlighting the complex regulatory landscape involving miRNAs and DEGs in DFUs. The identification of these hub genes and their associated miRNAs offers valuable insights into the molecular mechanisms driving DFU (43, 44).

In the context of diabetic foot ulcers (DFUs), several key genes and microRNAs have emerged as promising targets for therapeutic intervention due to their crucial roles in inflammation, cell survival, and tissue repair. TP53 (Tumor Protein p53), known for its role as the “guardian of the genome,” regulates the cell cycle, apoptosis, and DNA repair. Targeting TP53 through gene therapy or pharmacological agents could enhance cellular responses to stress and promote effective wound healing by restoring normal apoptotic processes and improving repair mechanisms. GAPDH (Glyceraldehyde-3-Phosphate Dehydrogenase), traditionally involved in glycolysis, also plays a role in cellular stress responses. Modulating GAPDH activity could improve energy metabolism and reduce chronic inflammation in DFUs, supporting the high metabolic demands of wound-healing cells. AKT1 (RAC-alpha Serine/Threonine-Protein Kinase) is a key player in cell survival, proliferation, and angiogenesis. Therapeutic strategies targeting AKT1 could involve developing inhibitors or activators to modulate cell survival and promote angiogenesis, enhancing blood supply to the wound site and facilitating tissue regeneration. MYC (Myc Proto-Oncogene Protein) regulates cell growth and metabolism. Therapeutic approaches could include using MYC inhibitors to control excessive proliferation and support metabolic processes that are critical for effective wound healing (45, 46). TNF (Tumor Necrosis Factor) is a central pro-inflammatory cytokine involved in chronic inflammation. Anti-TNF therapies could reduce persistent inflammation in DFUs, improving wound healing outcomes by targeting the inflammatory pathways that contribute to the chronicity of these ulcers. EGFR (Epidermal Growth Factor Receptor) is crucial for cell proliferation and migration. Therapeutic strategies could include EGFR inhibitors to regulate cell proliferation and growth factor therapies to enhance re-epithelialization and support wound closure. STAT3 (Signal Transducer and Activator of Transcription 3) modulates inflammation and cell survival. Inhibitors of STAT3 could reduce chronic inflammation and fibrosis, improving wound healing by enhancing cellular responses to injury. FN1 (Fibronectin 1) is essential for cell adhesion and ECM remodeling. Therapies targeting FN1 could include agents that enhance or mimic fibronectin activity to support cell migration and tissue repair in chronic wounds. VEGFA (Vascular Endothelial Growth Factor A) is a key factor in angiogenesis. VEGFA agonists or gene therapies could stimulate new blood vessel formation and improve tissue oxygenation, facilitating better wound healing. JUN (Transcription Factor Jun) regulates inflammation and cell proliferation. Therapeutic approaches could involve developing JUN inhibitors to reduce excessive inflammation and fibrosis, supporting effective tissue repair. MicroRNAs also offer valuable therapeutic potential. hsa-miR-34a-5p regulates apoptosis and the cell cycle; its modulation could restore normal apoptotic processes and improve wound healing. hsa-miR-155-5p is involved in inflammation, and targeting it could reduce chronic inflammatory responses in DFUs. hsa-miR-17-5p influences cell proliferation and angiogenesis, with potential therapies involving its modulation to support wound repair. hsa-miR-29b-3p regulates ECM remodeling and fibrosis, and targeting it could control fibrosis and enhance tissue regeneration. hsa-miR-7-5p, hsa-miR-1-3p, and hsa-miR-23b-3p are involved in regulating inflammation and cellular processes, offering additional targets for reducing inflammation and promoting wound healing. By leveraging the roles of these genes and microRNAs, novel therapeutic strategies can be developed to address the complex pathology of DFUs, ultimately improving patient outcomes and facilitating effective wound healing (47, 48).

The Gene Ontology (GO) analysis of diabetic foot ulcers (DFUs) highlights several critical areas for potential therapeutic intervention, focusing on biological processes, molecular functions, cellular components, and pathways. Among the biological processes, the Positive Regulation of miRNA Transcription (GO:1902895) is significantly enriched, indicating an upregulation of mechanisms that control microRNA (miRNA) transcription. Since miRNAs are crucial in regulating gene expression related to wound healing and inflammation, targeting these pathways could help modulate miRNA levels to enhance DFU management. Similarly, the Positive Regulation of miRNA Metabolic Process (GO:2000630) and Regulation of miRNA Transcription (GO:1902893) are also enriched, emphasizing the importance of controlling miRNA metabolism and overall transcription regulation to restore balance in miRNA expression and improve wound repair. In terms of molecular functions, the analysis identifies significant enrichment in DNA-binding Transcription Activator Activity (RNA Polymerase II-specific), which is essential for initiating the transcription of genes involved in inflammation and tissue repair. Modulating these transcriptional activities could influence gene expression related to DFU pathology. Additionally, the enrichment of functions like Protein Phosphatase Binding, Protease Binding, and General Transcription Initiation Factor Binding suggests that targeting these interactions could regulate cellular signaling pathways and improve cellular responses in DFUs. The analysis of cellular components reveals enrichment in Vesicle (GO:0031982) and Platelet Alpha Granule Lumen (GO:0031093), which are involved in cellular transport and storage of signaling molecules. Enhancing vesicle function could improve the delivery of growth factors and other critical molecules for wound healing. Other enriched components include the Nucleus and Intracellular Membrane-Bounded Organelle, which play roles in gene regulation and cellular organization. Targeting nuclear processes and membrane-bound organelles could influence cellular responses and support healing in DFUs. Finally, the analysis of pathways highlights significant enrichment in Proteoglycans in Cancer (p-value 4.696e-15), which are involved in cell signaling and extracellular matrix (ECM) remodeling. Targeting this pathway could address abnormal ECM interactions in DFUs. Additionally, pathways related to Human Cytomegalovirus Infection (p-values 2.422e-12, 2.772e-12) suggest a potential viral component in DFU pathology, pointing to the need for further investigation into viral infections’ role in DFUs. The enrichment of pathways like MAPK Signaling Pathway and Pathways in Cancer indicates that modulating these signaling pathways could influence inflammation and tissue repair, offering new therapeutic targets for managing DFUs. Overall, these insights from the GO analysis provide a comprehensive understanding of the underlying mechanisms in DFUs and suggest novel therapeutic strategies targeting miRNA regulation, molecular functions, cellular components, and specific pathways to improve wound healing and manage chronic inflammation (49–51).

## 5. Conclusion

In conclusion, the comprehensive analysis of diabetic foot ulcers (DFUs) highlights several critical targets for therapeutic intervention. The Gene Ontology (GO) analysis identifies significant enrichment in biological processes such as Positive Regulation of miRNA Transcription and Regulation of miRNA Transcription, underscoring the pivotal role of microRNAs like hsa-miR-34a-5p, hsa-miR-155-5p, hsa-miR-17-5p, hsa-miR-29b-3p, hsa-miR-7-5p, hsa-miR-1-3p, and hsa-miR-23b-3p in modulating gene expression related to wound healing and inflammation. Targeting these miRNAs could offer novel strategies for managing DFUs by restoring normal cellular functions and improving wound repair. The molecular functions enriched in the analysis, such as DNA-binding Transcription Activator Activity and Protein Phosphatase Binding, suggest that modulating key genes involved in these processes, including TP53, GAPDH, AKT1, MYC, TNF, EGFR, STAT3, FN1, VEGFA, and JUN, could improve gene regulation and cellular responses in DFUs. Additionally, the enriched cellular components, such as Vesicle and Platelet Alpha Granule Lumen, emphasize the importance of cellular transport and signaling. Targeting these components could enhance the delivery of critical molecules for wound healing. Pathways such as Proteoglycans in Cancer and Human Cytomegalovirus Infection reveal underlying mechanisms and potential viral influences in DFUs. Addressing these pathways could provide new avenues for treatment. By focusing on these enriched biological processes, molecular functions, and pathways, researchers can develop targeted therapies that address the complex pathology of DFUs, ultimately improving patient outcomes and advancing wound healing strategies.

## Notes

### Competing Interest Statement

The authors have declared no competing interest.

